# Sub-centrosomal mapping identifies augmin-γTuRC as part of a centriole-stabilizing scaffold

**DOI:** 10.1101/2020.11.18.384156

**Authors:** Nina Schweizer, Laurence Haren, Ricardo Viais, Cristina Lacasa, Ilaria Dutto, Andreas Merdes, Jens Lüders

## Abstract

Centriole biogenesis and maintenance are crucial for cells to generate cilia and assemble centrosomes that function as microtubule organizing centers (MTOCs). Centriole biogenesis and MTOC function both require the microtubule nucleator γ-tubulin ring complex (γTuRC). The widely accepted view is that γTuRC localizes to the pericentriolar material (PCM), where it nucleates microtubules. γTuRC has also been observed at centriolar regions that lack PCM, but the significance of these findings is unclear. Here we have used expansion microscopy to map spatially and functionally distinct sub-populations of centrosomal γTuRC including in the centriole lumen. Luminal localization is mediated by augmin and both complexes are linked to the centriole inner scaffold through POC5. Disruption of luminal localization impairs centriole stability and cilia assembly, defects that are also observed in γTuRC mutant fibroblasts derived from a patient suffering from microcephaly with chorioretinopathy. These results identify a novel, non-canonical role of augmin-γTuRC in the centriole lumen that is linked to human disease.

## Introduction

Centrioles, which are at the core of the centrosome and template the formation of cilia, are formed by nine sets of microtubules that are arranged in a circular fashion so that they form the wall of a cylinder. In human cells, wall microtubules of mature centrioles are organized as triplets and doublets in the proximal and distal cylinder, respectively. Triplets consist of one inner, complete microtubule, the so-called A-tubule, and two incomplete B- and C-tubules that share part of their wall with the adjacent A- and B-tubule, respectively. Doublets consist of only A- and B-tubules ^1^. In cycling cells formation of new centrioles is initiated during S phase and occurs laterally at mother centrioles ^2^. The origin of centriolar wall microtubules is not clear. The finding that the nucleator γTuRC is required for centriole biogenesis ^3–7^ and the observation that γTuRC-shaped structures cap the minus-end of A-tubules ^8^, suggest that at least the A-tubules arise by nucleation. During S/G2 phase daughter centrioles elongate and, after passing through mitosis, are converted to centrosomes through acquisition of PCM ^9^. The PCM is the canonical site of γTuRC localization, where it has a well-established role as a nucleator of microtubules that extend into the cytoplasm during interphase and are incorporated into the spindle during mitosis ^10–12^. Electron microscopy (EM) and, more recently, super resolution microscopy have revealed localization of γTuRC subunits also at the subdistal appendages ^13,14^ and in the lumen of mother centrioles ^5,15–17^, but their roles at these sites were not investigated. During ciliogenesis, the mother centriole is transformed into a basal body and templates formation of the axoneme, a microtubule-based scaffold structure that is at the core of cilia. However, axoneme microtubules are believed to not require nucleation but originate from elongation of the doublet microtubules in the distal basal body wall ^18^.

Here we have re-evaluated the long-standing view that γTuRC is a component of the PCM and that its centrosomal role is to nucleate microtubules. We found that γTuRC is distributed as functionally distinct sub-populations on the outside and, in complex with augmin, in the lumen of centrioles. Luminal augmin-γTuRC does not nucleate microtubules but contributes to centriole integrity, maintaining the ability of centrioles to template formation of cilia.

## Results

### γTuRC forms separable centrosomal sub-populations

To identify potential centrosomal sub-populations of γTuRC and elucidate whether these may have distinct functions, we analyzed the centrosomal localization of the γTuRC targeting factor NEDD1 and of the core subunits γ-tubulin and GCP4 by expansion microscopy (ExM). γTuRC subunits localized on the outer surface of both mother and daughter centrioles, visualized with anti-acetylated tubulin antibodies, in some cases displaying enrichment in the proximal part of mother centrioles. In addition, all three proteins were found in the centriole lumen (Fig. 1a,c,d,e). This localization pattern was fundamentally different from that of the bona-fide PCM components CDK5RAP2 and pericentrin, which associated only with the outer, proximal part of mother centrioles (Fig. 1a). To re-evaluate the paradigm that centrosomal microtubules are nucleated in the PCM, we analyzed microtubule regrowth after cold-induced depolymerization. Whereas microtubules could not be detected in cold-treated cells, after a few seconds of warming microtubules were nucleated in close proximity of centriole cylinders (Fig. 1b). Microtubules grew preferentially from the proximal surface of mother centrioles, but were also observed along the entire centriole wall including at distal ends. Thus, γTuRC and nucleation activity are generally associated with the outer surface of centrioles, with some enrichment in the region of the PCM.

**Figure 1.**
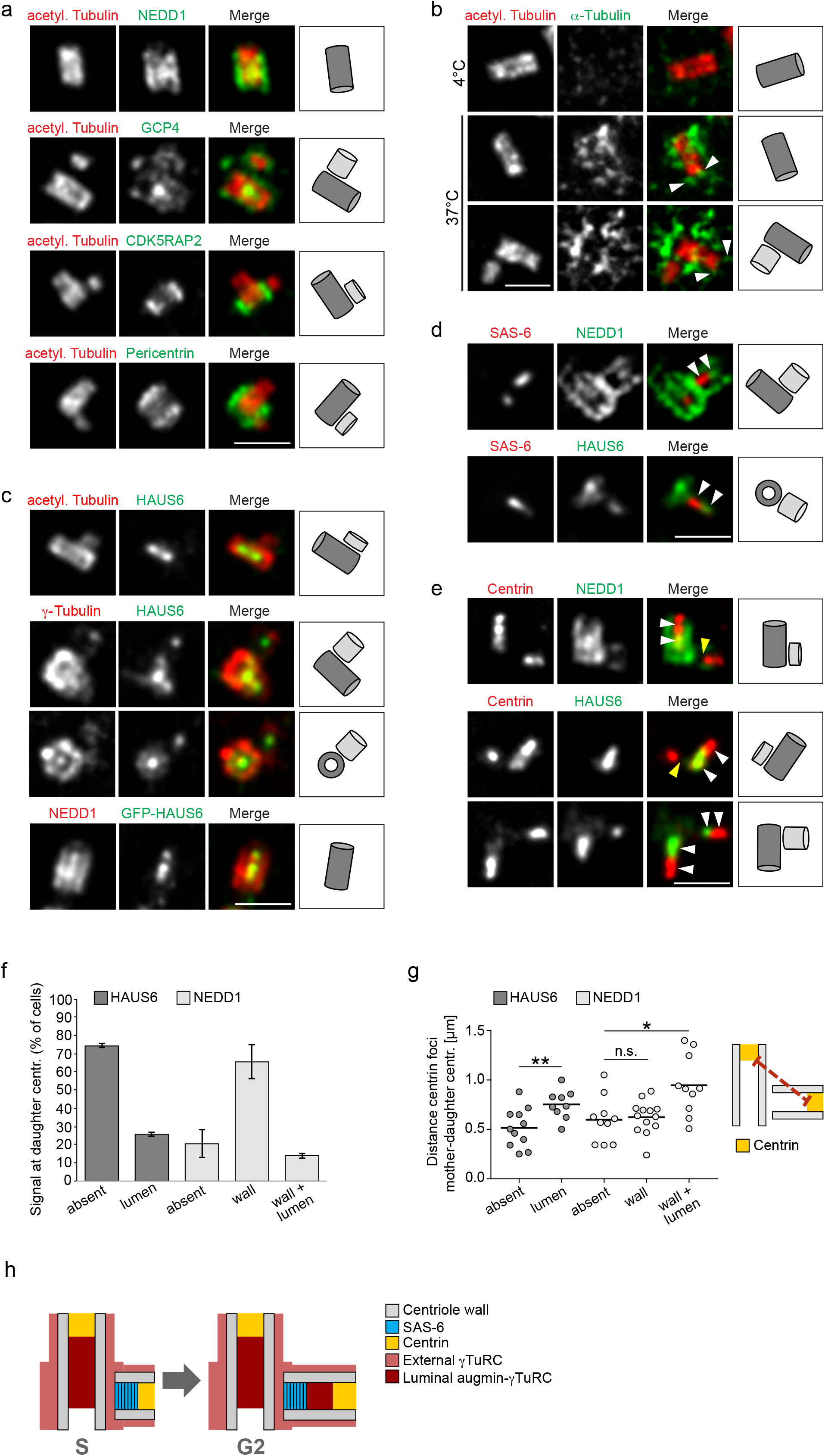
γTuRC forms distinct centrosomal sub-populations. (**a**) Centrioles of U2OS cells in ExM stained for acetylated α-tubulin (red) and either NEDD1 (green), GCP4 (green), CDK5RAP2 (green) or pericentrin (green). (**b**) Centrioles and microtubules of interphase U2OS cells in ExM stained for acetylated α-tubulin (red) and α-tubulin (green) in a microtubule regrowth assay. Depicted is the condition before (4°C) and after (37°C) microtubule regrowth. Arrowheads point to microtubules associated with the distal centriole wall. (**c**) Centrioles of parental U2OS cells or U2OS cells stably expressing EGFP-HAUS6 in ExM stained for acetylated α-tubulin (red) and HAUS6 (green), γ-tubulin (red) and HAUS6 (green) or NEDD1 (red) and GFP (EGFP-HAUS6, green). (**d**) Centrioles of U2OS cells in ExM stained for SAS-6 (red) and either NEDD1 (green) or HAUS6 (green). Arrowheads point to adjacent signals of NEDD1/SAS-6 or HAUS6/SAS-6 in the daughter centriole lumen. (**e**) Centrioles of U2OS cells in ExM stained for centrin (red) and either NEDD1 (green) or HAUS6 (green). White arrowheads point to adjacent signals of NEDD1/centrin or HAUS6/centrin in the lumen of mother and daughter centrioles. Yellow arrowheads point to daughter centrioles that lack NEDD1 or HAUS6 in the lumen. (**f**) Quantifications of the percentage of cells with HAUS6 or NEDD1 at the wall/lumen of daughter centrioles. Error bars represent standard deviations from the mean obtained from two independent experiments (HAUS6 (absent): 74.3 ± 1.1%, HAUS6 (lumen): 25.8 ± 1.1%, NEDD1 (absent): 20.5 ± 7.8%, NEDD1 (wall): 65.5 ± 9.2%, NEDD1 (wall and lumen): 14.0 ± 1.4%, mean ± SD, 100-102 cells per condition and experiment). (**g**) Quantifications of the distance between the centrin foci of mother and daughter centrioles in cells where HAUS6 or NEDD1 are absent/present at daughter centrioles. One experiment, means are depicted as horizontal lines (HAUS6 (absent): 0.5 ± 0.2 μm, HAUS6 (lumen): 0.8 ± 0.2 μm, NEDD1 (absent): 0.6 ± 0.2 μm, NEDD1 (wall): 0.6 ± 0.2 μm, NEDD1 (wall and lumen): 0.9 ± 0.3 μm, mean ± SD, 9-13 mother-daughter centriole pairs from 9-13 cells per condition, p (HAUS6 absent/lumen) < 0.01, p (NEDD1 absent/wall) = 0.762, p (NEDD1 absent/wall and lumen) < 0.05). (**h**) Cartoon summarizing the localizations of distinct γTuRC sub-populations. Cartoons in (**a**-**e**) illustrate centriole configurations in the corresponding panels, dark grey = mother centriole, light grey = daughter centriole. Bar (all panels), 2 μm. ** p < 0.01, * p < 0.05, n.s. = not significant

### γTuRC co-localizes with augmin in the central 92 lumen of centrioles

Next, we focused on luminal γTuRC. Interestingly, we found that subunits of the augmin complex, which recruits γTuRC to spindle microtubules during mitosis ^19,20^, colocalized with γTuRC in the centriole lumen during interphase (Fig. 1c). Luminal localization was observed for both endogenous augmin subunits and EGFP-tagged recombinant versions (Fig. 1c, Supplementary Fig. 1a). By comparing the localizations of HAUS6 and NEDD1 relative to SAS-6, which marks the cartwheel structure at the proximal end of daughter centrioles ^21^, and centrin, a marker for the distal lumen of centrioles ^22^, we found that in all cases NEDD1 and HAUS6 were found distal to the SAS-6 signal (Fig. 1d) and proximal to the bulk of the centrin signal (Fig. 1e). These results show that apart from γTuRC that is recruited to the outside of centrioles, a separate pool colocalizes with augmin in the central region of the centriole lumen.

### Lumen recruitment of augmin and γTuRC occurs late during centriole elongation

Interestingly, newly formed daughter centrioles frequently lacked luminal augmin and γTuRC, even though γTuRC was visible on the outside of the daughter cylinder (Fig. 1e). Consistent with this, only a minor fraction of all daughter centrioles identified by centrin labeling, was also positive for luminal HAUS6 or NEDD1 staining (Fig. 1f). Measuring the distance between centrin foci of mother and daughter centrioles as a proxy for daughter centriole length revealed that these centrioles were more elongated than daughter centrioles that lacked these proteins (Fig. 1g), suggesting that augmin and γTuRC localize to the lumen late during centriole biogenesis. Corroborating this result, accumulation of HAUS6 in the lumen of daughters coincided with poly-glutamylation, a tubulin modification that occurs selectively on the C-tubules of triplet microtubules ^23^ and that became detectable only after daughters had reached a substantial length (Supplementary Fig. 1b). The most robust luminal HAUS6 signal was observed in mature centrioles, where it was confined to the proximal/central cylinder, marked by poly-glutamylation (Supplementary Fig. 1b).

Together our results indicate that γTuRC localizes first to the outside of newly formed daughter centrioles and subsequently, during centriole elongation, accumulates with augmin in the centriole lumen (Fig. 1h).

### Centriole outer wall recruitment of γTuRC depends on CEP192

Previous work implicated CEP192 in the recruitment of γTuRC to centrosomes ^24,25^ and identified the targeting factor NEDD1 as proximity interactor of CEP192 ^26,27^, but did not distinguish between distinct sub-centrosomal sites. Super resolution microscopy detected CEP192 along the outer walls of mother and daughter centrioles (Fig. 2b) ^28^. To re-evaluate CEP192’s role in γTuRC centrosome recruitment, cells were transfected with siRNA, synchronized in mitosis with the Eg5 inhibitor STLC and then, bypassing cell division, released into G1 by CDK1 inhibition (Fig. 2a). This setup avoided adverse effects on centrosomes from duplication failure ^24^, and enriched interphase cells with fully elongated, CEP192-depleted centrioles. We found that CEP192 depletion removed NEDD1 specifically from the outside of centrioles, whereas the luminal pool of NEDD1 appeared unaffected (Fig. 2b,c).

**Figure 2.**
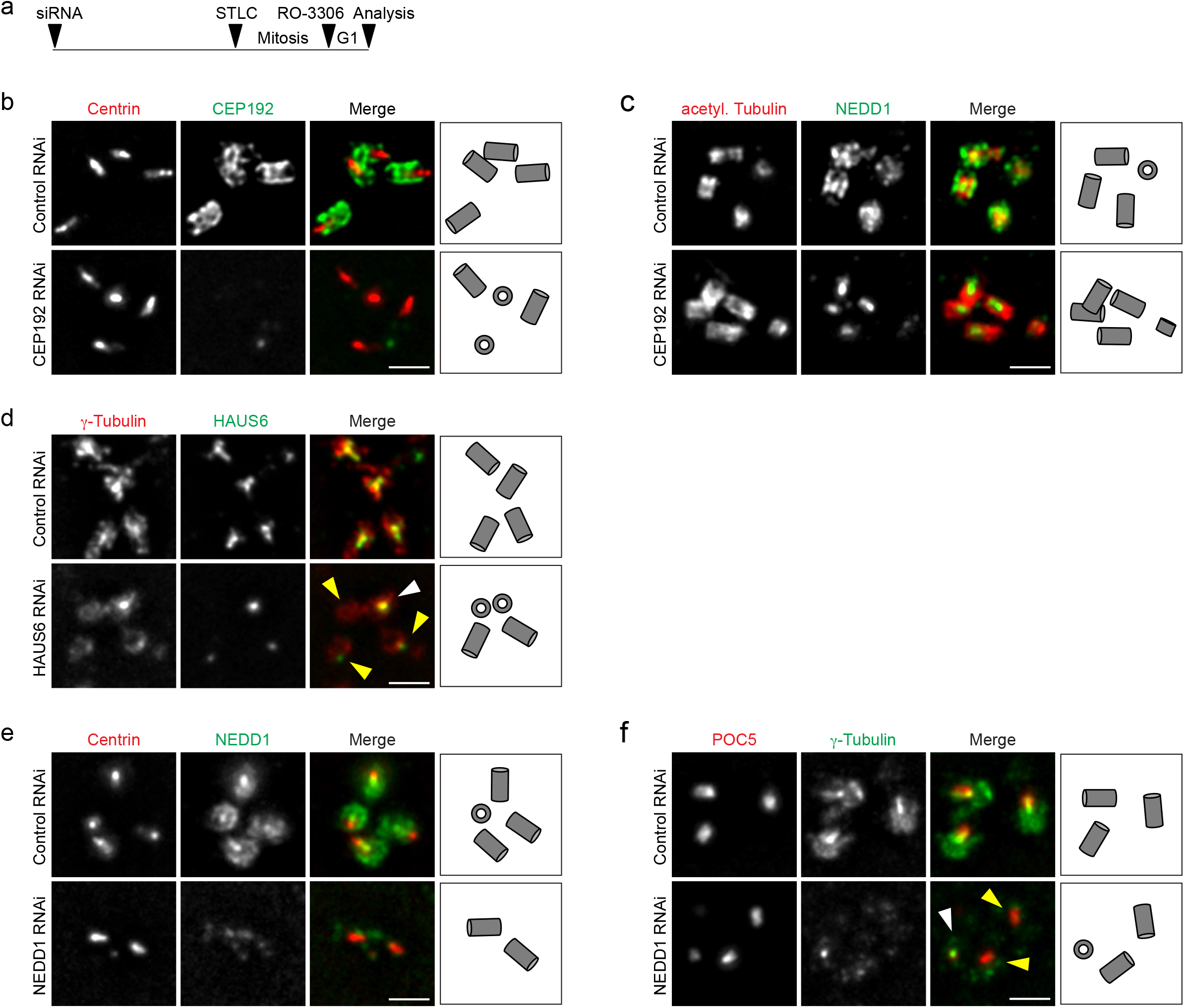
γTuRC centriole localization depends on CEP192 and augmin. (**a**) Schematic depicting the experimental design. ~52 h after siRNA transfection, STLC was added to the culture medium for ~18 h to arrest cells in mitosis. Mitotic cells were collected, released into G1 by addition of RO-3306 and fixed ~4 h later. (**b**) Centrioles of control RNAi and CEP192 RNAi U2OS cells in ExM stained for centrin (red) and CEP192 (green). (**c**) Centrioles of control RNAi and CEP192 RNAi U2OS cells in ExM stained for acetylated α-tubulin (red) and NEDD1 (green). (**d**) Centrioles of control RNAi and HAUS6 RNAi U2OS cells in ExM stained for γ-tubulin (red) and HAUS6 (green). White arrowhead points to a centriole that is not depleted of γ-tubulin, yellow arrowheads point to centrioles that lack HAUS6/luminal γ-tubulin. (**e**) Centrioles of control RNAi and NEDD1 RNAi U2OS cells in ExM stained for centrin (red) and NEDD1 (green). (**f**) Centrioles of control RNAi and NEDD1 RNAi U2OS cells in ExM stained for POC5 (red) and γ-tubulin (green). White arrowhead points to a centriole that has γ-tubulin in the lumen, yellow arrowheads point to centrioles that are depleted of γ-tubulin at the wall and in the lumen. Bar (all panels), 2 μm. Cartoons illustrate centriole configurations in the corresponding panels.

### Lumen recruitment of γTuRC requires augmin

Considering their colocalization (Fig. 1c), we asked whether luminal recruitment of γTuRC required augmin. Using the same experimental setup as before, we depleted cells of HAUS6 (Supplementary Fig. 2a). Strikingly, centrioles lacking HAUS6 also lacked γ-tubulin in the lumen (Fig. 2d), whereas γ-tubulin signals on the outside of centrioles could still be detected (Fig. 2d). Importantly, we never observed HAUS6-negative centrioles that were positive for luminal γ-tubulin. We additionally analyzed γTuRC lumen localization in mouse hippocampal neurons in which *Haus6* had been knocked out post-mitotically. We previously showed that during neuronal culture PCM-associated γ-tubulin is strongly downregulated and the remaining signal is centriole-associated ^29^. Indeed, in neurons at nine days in vitro (DIV) both HAUS6 and γ-tubulin displayed centriolar localization. Strikingly, both proteins were largely absent from centrioles in conditional *Haus6* KO neurons (Supplementary Fig. 2b,c), indicating that residual centrosomal γTuRC in differentiated neurons is luminal and recruited by the same augmin-dependent mechanism as in cycling cells. Since previous work showed that γTuRC targeting is broadly mediated by NEDD1 ^3,4^, we also tested depletion of NEDD1. In this condition NEDD1 and γ-tubulin were absent from both the outside and the lumen of centrioles (Fig. 2e,f). Thus, outer wall localization of γTuRC requires CEP192, luminal γTuRC localization depends on augmin, and the targeting factor NEDD1 is required for recruitment to both sites.

### Centriolar augmin interacts with the inner scaffold protein POC5

To learn more about the roles of augmin and γTuRC in the centriole lumen, we performed biotin proximity labeling using HAUS6 fused to the BirA biotin ligase as bait and identified centriole-specific, biotinylated proteins by mass spectrometry (Fig. 3a, Supplementary Fig. 3a,b). This approach identified POC5, a centrin-binding protein and component of a scaffold structure at the luminal surface of centrioles proposed to protect against mechanical stress ^30–32^. Consistent with this, augmin subunits were previously found as proximity interactors in cells expressing POC5-BirA as bait ^26^. POC5 was present in the lumen of mother centrioles and accumulated in the lumen of daughter centrioles after these had reached a significant length (Fig. 3b), resembling the behavior that we had observed for luminal augmin and γTuRC. POC5 and γ-tubulin were confined to the same central luminal region, but in end-on views γ-tubulin appeared to localize more interior than POC5 (Fig. 3b).

**Figure 3.**
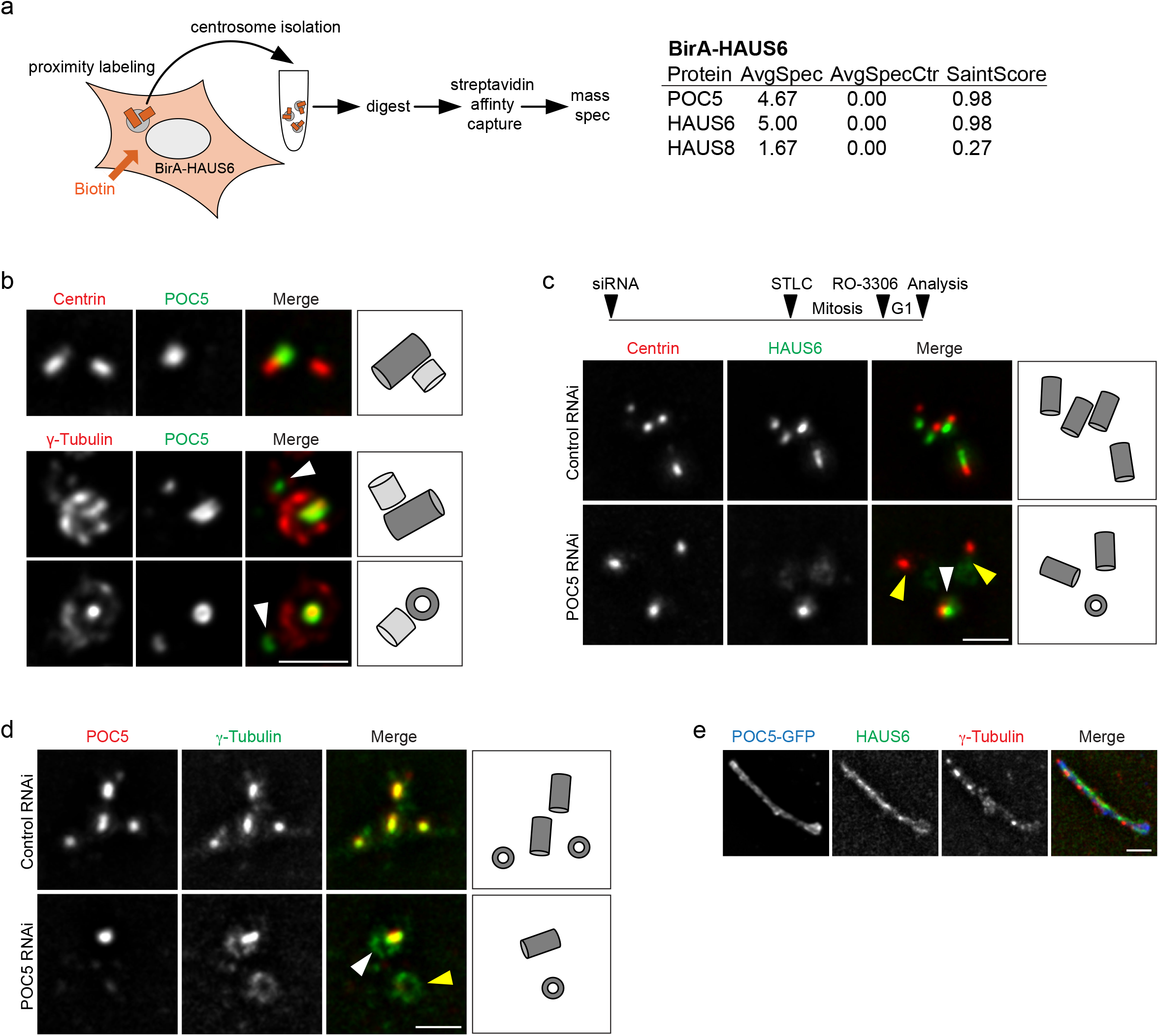
Augmin is recruited to the inner centriole scaffold by POC5. (**a**) Mass spectrometry analysis of proximity interactors of BirA-HAUS6 from isolated centrosomes of U2OS cells. AvgSpec = average spectral counts, AvgSpecCtrl = average spectral counts in the control (parental U2OS cells). (**b**) Centrioles of U2OS cells in ExM stained for centrin (red) and POC5 (green) or γ-tubulin (red) and POC5 (green). Arrowheads point to POC5 in the daughter centriole lumen. (**c,d**) Centrioles of control RNAi and POC5 RNAi U2OS cells in ExM stained for centrin (red) and HAUS6 (green) or POC5 (red) and γ-tubulin (green). White arrowheads point to centrioles with HAUS6 or γ-tubulin in the lumen, yellow arrowheads point to centrioles that lack HAUS6 or luminal γ-tubulin. Bar, 2 μm. The schematic in (**c**) depicts the experimental design in (**c,d**). ~52 h after siRNA transfection, STLC was added to the culture medium for ~18 h to arrest cells in mitosis. Mitotic cells were collected, released into G1 by addition of RO-3306 and fixed ~4 later. (**e**) POC5-GFP aggregate in ExM stained for GFP (POC5-GFP, blue), HAUS6 (green) and γ-tubulin (red). Bar (all panels), 2 μm. Cartoons in (**b**,**c**,**d**) illustrate centriole configurations in the corresponding panels, dark grey = mother centriole, light grey = daughter centriole.

### POC5 is required for luminal recruitment of augmin-γTuRC

Since we occasionally observed daughter centrioles that were positive for POC5, but lacked γ-tubulin in the lumen (Fig. 3b), we tested whether POC5 may function upstream of augmin and γTuRC lumen recruitment. Using the mitotic arrest-release approach, we found that centrioles depleted of POC5 also lacked luminal HAUS6 and γ-tubulin, whereas both proteins were always present at centrioles in control cells (Fig. 3c,d). Previous analysis by cryo-electron tomography (cryo-ET) showed that the inner scaffold is a periodic, helical structure, lining the inner centriole wall ^32^, likely composed of repeating units of scaffold protein complexes. Thus, scaffold proteins may tend to self-associate. Indeed, POC5 exogenously expressed in human cells forms filamentous structures in the cytoplasm that associate with other centriole proteins ^30^. We confirmed this observation and found that these ectopic assemblies were also labeled with antibodies against HAUS6 and γ-tubulin (Fig. 3e, Supplementary Fig. 3d). Together, these findings demonstrate that POC5, through interaction with augmin, recruits γTuRC to the inner centriole scaffold.

### POC5 and augmin promote centriole integrity

The inner scaffold was suggested to confer stability on centrioles ^32^. When we quantified the number of centrioles at the end of the duplication cycle, by counting of centrin foci in mitotic cells, we found that POC5 depletion had no effect on the number of centrioles (Supplementary Fig. 4a). We also did not observe any change in centriole number after HAUS6 depletion (Supplementary Fig. 4b). However, when mitotic duration was extended up to ~18 h by treatment with the Eg5 inhibitor STLC, centriole numbers in POC5 and HAUS6 RNAi cells declined: whereas 70-80% of control cells still had the expected number of at least 4 centrin foci, this number was observed in only ~35% of POC5 depleted cells and ~50% of HAUS6 depleted cells (Fig. 4a,b,c,d), suggesting centriole destabilization. Time course experiments further revealed that the decline in centriole numbers correlated with the time spent in mitosis (Fig. 4e). We also assayed centriole stability in the non-cancer RPE1 cell line. To avoid p53-dependent G1 arrest caused by mitotic defects ^33^, we induced cell cycle exit by serum withdrawal immediately after transfection of siRNA. After HAUS6 was efficiently depleted, we added serum for cell cycle re-entry, and STLC for prolonged mitotic arrest. Similar to U2OS cells, albeit less pronounced, centrioles in HAUS6-depleted RPE1 cells were also destabilized (Supplementary Fig. 4c).

**Figure 4.**
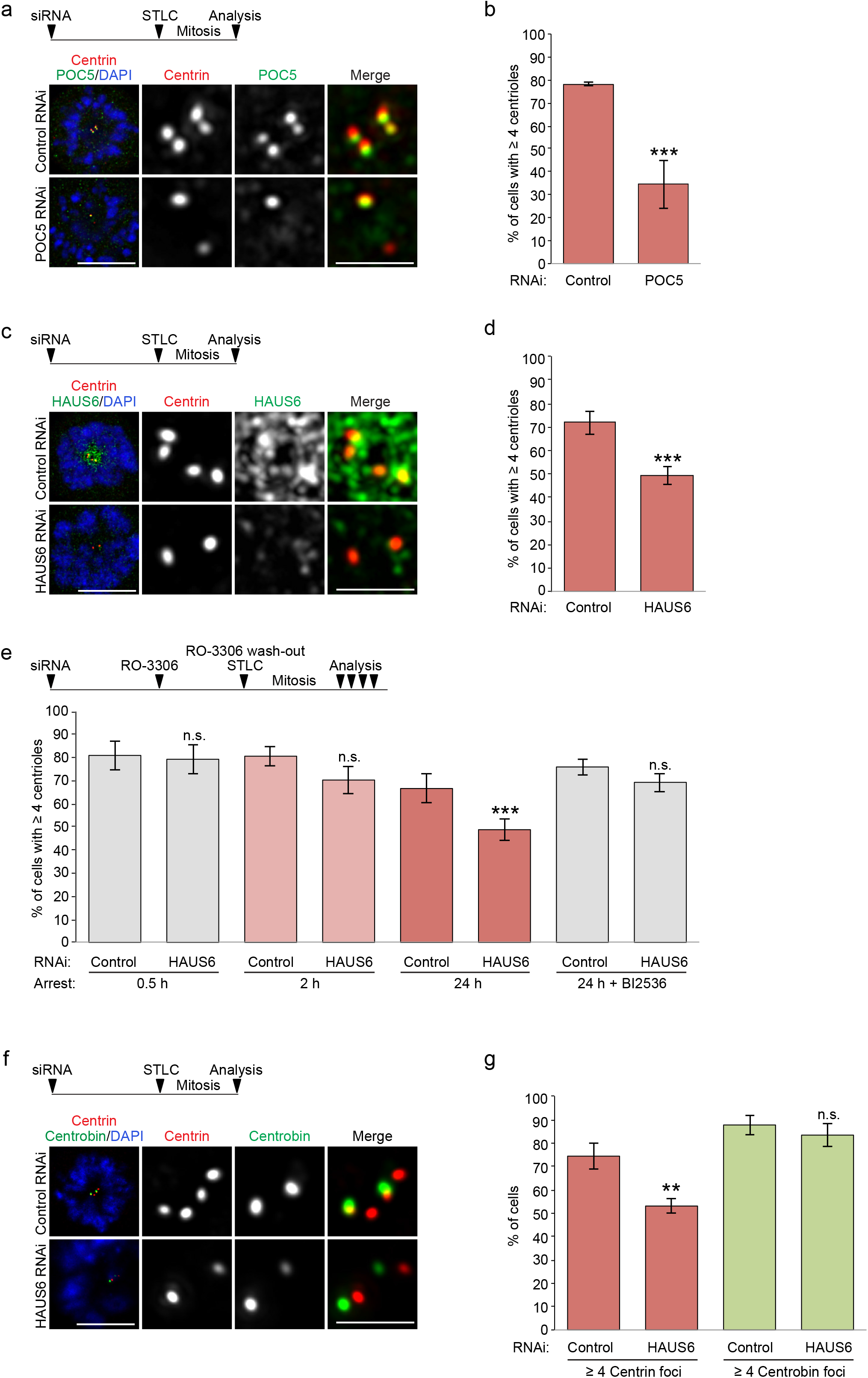
POC5 and augmin promote centriole stability. (**a**) Control RNAi and POC5 RNAi U2OS cells arrested in mitosis with STLC stained for centrin (red), POC5 (green) and DNA (DAPI, blue). The experimental design is depicted schematically. ~52 h after siRNA transfection, STLC was added to the culture medium for ~18 h. (**b**) Quantifications of the percentage of control RNAi or POC5 RNAi U2OS cells with ≥ 4 centrioles (centrin foci) after spending up to ~18 h in mitosis. Error bars represent standard deviations from the mean obtained from three independent experiments (control RNAi: 78 ± 0.7%, POC5 RNAi: 34.7 ± 10.6%, mean ± SD, 239-337 cells per condition and experiment, p < 0.001). Control RNAi and HAUS6 RNAi U2OS cells arrested in mitosis with STLC stained for centrin (red), HAUS6 (green) and DNA (DAPI, blue). The experimental design is depicted schematically. ~52 h after siRNA transfection, STLC was added to the culture medium for ~18 h. (**d**) Quantifications of the percentage of control RNAi or HAUS6 RNAi U2OS cells with ≥ 4 centrioles (centrin foci) after spending up to ~18 h in mitosis. Error bars represent standard deviations from the mean obtained from four independent experiments (control RNAi: 71.8 ± 4.9%, HAUS6 RNAi: 49.5 ± 3.9%, mean ± SD, 232-335 cells per condition and experiment, p < 0.001). (**e**) Quantifications of the percentage of control RNAi or HAUS6 RNAi U2OS cells with ≥ 4 centrioles (centrin foci) after spending different times in mitosis, in the presence or absence of PLK1 inhibitor BI2536. Error bars represent standard deviations from the mean obtained from three to six independent experiments (control RNAi (0.5 h): 81 ± 6.2%, HAUS6 RNAi (0.5 h): 79.3 ± 6.5%; mean ± SD, 3 experiments, 154-329 cells per condition and experiment, p = 0.765; control RNAi (2 h): 80.7 ± 4.2%, HAUS6 RNAi (2 h): 70.7 ± 5.5%, mean ± SD, three experiments, 125-250 cells per condition and experiment, p = 0.071; control (24 h): 66.7 ± 6.4%, HAUS6 RNAi (24 h): 48.8 ± 4.6%; mean ± SD, 6 experiments, 115-350 cells per condition and experiment, p < 0.001; control RNAi (24 h, BI2536): 76.0 ± 3.5%, HAUS6 RNAi (24 h, BI2536): 69.3 ± 4.0%; mean ± SD, 3 experiments, 110-311 cells per condition and experiment, p = 0.100). The experimental design is depicted schematically. ~52 h after siRNA transfection, RO-3306 was added to the culture medium for ~18 h to arrest cells in G2. RO-3306 was washed out and STLC was added to arrest cells in mitosis for defined time points. In some cases, BI2536 was added together with STLC, as indicated. (**f**) Control RNAi and HAUS6 RNAi U2OS cells arrested in mitosis with STLC stained for centrin (red), centrobin (green) and DNA (DAPI, blue). The experimental design is depicted schematically. ~52 h after siRNA transfection, STLC was added to the culture medium for ~18 h. (**g**) Quantifications of the percentage of control RNAi or HAUS6 RNAi U2OS cells with ≥ 4 centrin foci or ≥ 2 centrobin foci after spending up to ~18 h in mitosis. Error bars represent standard deviations from the mean obtained from three independent experiments (% of cells with ≥ 4 centrin foci, control RNAi: 74.3 ± 5.5%, HAUS6 RNAi: 53.0 ± 3.0%, p < 0.01; % of cells with ≥ 2 centrobin foci, control RNAi: 87.7 ± 4.2%, HAUS6 RNAi: 83.3 ± 5.1%; mean ± SD, 300-504 cells per condition and experiment, p = 0.32). *** p < 0.001, ** p < 0.01; n.s., not significant. Bar (all panels), 10 μm (merge with DAPI) or 2 μm (insets depicting centrioles).

Curiously, during prolonged mitotic arrest the number of centrioles in control cells also slightly decreased (Fig 4e). We speculated that this was due to premature loss of the cartwheel, which normally occurs at mitotic exit in a PLK1-dependent manner ^9,34^. Indeed, in the presence of the PLK1 inhibitor BI2536 the cartwheel component SAS-6 was retained at daughter centrioles during prolonged mitotic arrest (Supplementary Fig. 4d,e). Counting of centrioles in PLK1-inhibited cells after prolonged mitotic arrest revealed that there was no significant difference in the percentage of cells with 4 or more centrioles between control and HAUS6 RNAi samples (~76% and ~69%; Fig. 4e), suggesting that centriole destabilization specifically affected cartwheel-less daughter centrioles.

The time-dependent disappearance of centrin foci in mitotically arrested cells may indicate complete centriole disassembly or merely loss of their distal, centrin-containing compartment. To distinguish between these possibilities, we quantified foci of centrobin, which localizes to the outer wall of daughter centrioles, in a central region ^35,36^. In contrast to the reduction in centrin foci, a similar percentage of control and HAUS6 depleted cells had the expected number of at least two centrobin foci (~88% versus ~83%) (Fig. 4f,g), demonstrating that centrioles did not completely disassemble.

### POC5 and γTuRC are required for ciliogenesis

While centriole destabilization was only observed upon prolonged mitosis, we hypothesized that cilia assembly, which relies on the elongation of microtubule doublets at the distal tip of mother centrioles, might naturally be sensitive to centriolar defects caused by the absence of luminal augmin-γTuRC. Since we were not able to efficiently remove HAUS6 from mother centrioles, we tested depletion of POC5, the scaffold protein most proximal to luminal HAUS6 (Fig. 3a). RPE1 cells were treated with control or POC5 siRNA and serum-starved to induce ciliogenesis. Whereas 67% of control cells had a cilium, only ~30% of POC5-depleted cells that lacked POC5 on both centrioles were ciliated (Fig. 5a,b). Since POC5 depletion was shown to cause G1 arrest in RPE1 cells ^31^, which could interfere with assaying ciliogenesis, we repeated the experiment in RPE1 *p53* KO cells. In this case ~55% of control cells were ciliated, whereas only ~14% of cells that lacked POC5 on both centrioles possessed a cilium (Fig. 5b), confirming that loss of POC5 from the centriole lumen impairs ciliogenesis.

**Figure 5.**
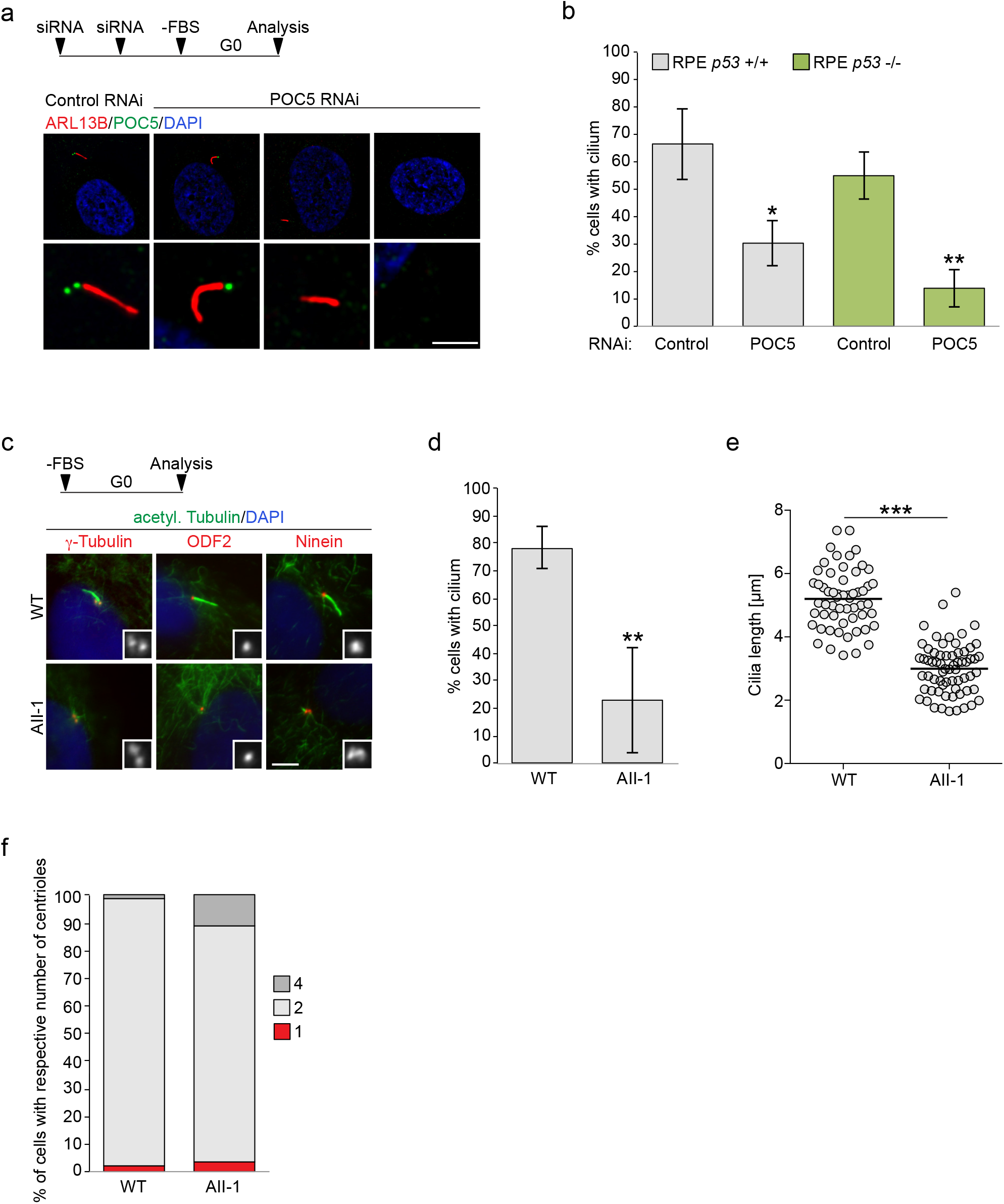
POC5 and γTuRC are required for ciliogenesis. (**a**) Serum-starved control RNAi and POC5 RNAi hTERT RPE1 cells stained for ARL13B (red), POC5 (green) and DNA (DAPI, blue). Bar, 10 μm (merge with DAPI) or 2 μm (insets depicting centrioles). The experimental design is depicted schematically. Cells were transfected two times with siRNA (second transfection after ~48 h) and serum-starved after a total of ~80-96 h for 48 h before cells were fixed. (**b**) Quantifications of the percentage of ciliated control RNAi and POC5 RNAi hTERT RPE1 and hTERT RPE1 *p53 -/-* cells. Error bars represent standard deviations from the mean obtained from three independent experiments (control RNAi hTERT RPE1: 66.6 ± 13.1%, POC5 RNAi hTERT RPE1: 30.4 ± 8.4%, mean ± SD, 118-202 cells per condition and experiment, p < 0.05; control RNAi hTERT RPE1 *p53* -/-: 55.0 ± 8.8%, POC5 RNAi hTERT RPE1 *p53* -/-: 13.9 ± 6.9%, mean ± SD, 82-113 cells per condition and experiment; p < 0.01). (**c**) Serum-starved human fibroblasts, obtained from a control individual (WT) or a patient with a mutation in GCP4 (AII-1) stained for acetylated α-tubulin (green) and either γ-tubulin (red), ODF2 (red) or ninein (red) and DNA (DAPI, blue). Insets show γ-tubulin, ODF2 or ninein, respectively. Bar, 5 μm. (**d**) Quantifications of the percentage of ciliated control (WT) or patient (AII-1) fibroblasts. Error bars represent standard deviations from the mean obtained from three independent experiments (WT: 78.3% ± 7.5%, AII-1: 23.0 ± 19.0%, mean ± SD, 99-122 cells per condition and experiment; p < 0.01). (**e**) Quantifications of cilia length in serum-starved control (WT) or patient (AII-1) fibroblasts. The mean is depicted as horizontal line (control: 5.2 ± 0.9 μm, AII-1: 3.0 ± 0.8 μm, mean ± SD, n = 57-65 cilia per condition, combined from three independent experiments, p < 0.001). (**f**) Quantifications of the percentage of serum-starved control (WT) or patient (AII-1) fibroblasts with 1, 2 or 4 centrioles. One experiment (WT (1 centriole): 1.9%, AII-1 (1 centriole): 3.5%, WT (2 centrioles): 97.1%, AII-1 (2 centrioles): 85.2%; WT (4 centrioles): 1.0%, AII-1 (4 centrioles): 11.3%, n (WT) = 105 cells, n (AII-1) = 142 cells. Schematics depict the design of experiments. *** p < 0.001, ** p < 0.01; * p < 0.05.

Mutations in γTuRC subunits have been linked to developmental defects including primary microcephaly and retinopathy ^37–41^. We hypothesized that some of the clinical manifestations may also involve centriole destabilization and impaired ciliogenesis. To address this, we analyzed the ability of GCP4 mutant fibroblasts, obtained from a patient diagnosed with microcephaly and chorioretinopathy^38^, to assemble cilia after serum starvation. Strikingly, only ~20% of GCP4 mutant fibroblasts were ciliated compared to ~80% of control fibroblasts (Fig. 5c,d). Additionally, cilia in patient fibroblasts were significantly shorter (Fig. 5e). Importantly, impaired ciliogenesis was not the consequence of defective centriole duplication (Fig. 5f). Moreover, mother centrioles had acquired subdistal appendages, as determined by the presence of ODF2 and ninein (Fig. 5c), indicating proper maturation. Thus, in GCP4 mutant cells centrioles form and mature, but are defective in supporting cilia assembly and growth.

## Discussion

Here we have identified novel, non-canonical roles of augmin and γTuRC in the centriole lumen that are independent of their previously described functions in microtubule nucleation. In addition to nucleating microtubules, both protein complexes are linked to the centriole inner scaffold through POC5, contributing to centriole integrity. POC5 was recently mapped to the innermost region of the scaffold, a position well suited for anchoring augmin-γTuRC ^32^. Augmin is known to directly interact with microtubules through its HAUS8 subunit ^42^ and this interaction is independent of its ability to recruit γTuRC ^42,43^. Thus, it is tempting to speculate that augmin, apart from interacting with POC5, may also directly bind and stabilize microtubules of the centriole wall. Curiously, shape and dimension of native and reconstituted augmin ^42,43^ have striking similarity to unassigned Y- and L-shaped linker structures that are connected to A- and B-tubules of the centriole wall and protrude into the lumen ^44–46^. However, further work is needed to more precisely map the configuration of luminal augmin.

Considering the periodicity of the inner scaffold structure ^32,44^ and our observation that augmin and γTuRC are distributed along the entire length of the central lumen, one can speculate that multiple copies of augmin and γTuRC may adopt a stacked configuration at the inner centriole wall. Since both augmin and γTuRC are large, multi-subunit complexes, they would be expected to form prominent structures in the centriole lumen. Indeed, several EM and cryo-ET studies have described unidentified densities including ring-shaped structures in the centriole lumen ^44,45,47,48^. The fact that these are not consistently observed may indicate sensitivity of augmin and γTuRC to the conditions used for sample preparation. While the precise arrangement of luminal augmin and γTuRC remains to be determined, we showed that their specific loss from the lumen impairs centriole integrity and ciliogenesis, similar to defects observed after depletion of WDR90, which functions more upstream in the recruitment of scaffold proteins ^46^. Based on our results, augmin and γTuRC may be considered components of the inner scaffold, which may thus extend farther into the centriole lumen than previously anticipated. Our findings also suggest that the integrity of the entire extended scaffold structure is required for its centriole-stabilizing and ciliogenesis-promoting function.

Importantly, the impairment of ciliogenesis caused by displacement of luminal augmin-γTuRC is also observed in patient-derived γTuRC mutant fibroblasts. Thus, phenotypes previously ascribed to augmin or γTuRC deficiency may not solely be related to their function as microtubule nucleators, but also to their luminal roles in promoting centriole integrity.

## Methods

### Cell culture and treatments

U2OS cells and hTERT RPE1 cells were cultured in DMEM and DMEM/F12 (Invitrogen), respectively, with 10% fetal bovine serum (FBS) and PenStrep (both Gibco). Parental cell lines were obtained from ATCC, hTERT RPE1 *p53* -/- cells were provided by Meng-Fu Bryan Tsou ^49^. U2OS cells stably expressing POC5-GFP, EGFP-HAUS8, EGFP-HAUS6 or BirA-HAUS6 were generated by transfection of the appropriate expression plasmid, followed by either the selection with 1 mg/ml geneticin (Gibco) or by FACS. Human fibroblasts were derived from skin biopsis from a control individual (WT) and a patient with GCP4 mutations (AII-1) ^38^ and cultured in DMEM with 15% FBS and PenStrep. All cell lines were kept in a 37°C incubator with 5% CO_2_ and a humidified atmosphere. For the induction of ciliogenesis, cells were incubated in DMEM without FBS for 48 h. G2 arrest or mitotic exit was induced with 10 μM RO-3306 (Sigma). Mitotic arrest was induced with 10 μM STLC (Sigma). PLK1 was inhibited with 100 nM BI2536 (Adooq Bioscience).

### Generation of mouse strains

A neuronal specific *Haus6* conditional KO mouse strain was generated by crossing *Haus6* floxed (*Haus6*^fl^) mice ^50^ (RBRC09630, Accession No. CDB1354K (http://www2.clst.riken.jp/arg/mutant%20mice%20list.html) with B6.Tg(Actl6b-Cre)4092Jiwu/J mice (Jackson Laboratories). Mouse strains were maintained on a mixed C57BL/6 background in strict accordance with the European Community (2010/63/UE) guidelines in the Specific-Pathogen Free (SPF) animal facilities of the Barcelona Science Park (PCB). All protocols were approved by the Animal Care and Use Committee of the PCB/University of Barcelona (IACUC; CEEA-PCB) and by the Departament de Territori I Sostenibilitat of the Generalitat de Catalunya in accordance with applicable legislation (Real Decreto 53/2013).

### Mice genotyping

DNA was extracted from tail biopsies by digesting biopsies with 0.4 mg/mL Proteinase K in 10 mM Tris-HCl, 20 mM NaCl, 0.2% SDS and 0.5 mM EDTA overnight at 56°C, followed by DNA precipitation with isopropanol. Genotyping was performed by PCR using the following primers: mAug6KO_FW, 5’-CAACCCGAGCAACAGAAACC-3’ and mAug6KO_Rev, 5’-CCTCCCACCAACTACAGACC-3’ to detect *Haus6* WT, *Haus6* floxed and *Haus6* KO alleles; 26994, 5’-GCTGGAAGATGGCGATTAGC-3’ and 30672, 5’-TCAGCCTGGTTACAAGGAACA-3’ to detect the Cre-recombinase transgene, primers olMR7338, 5’-CTAGGCCACAGAATTGAAAGATCT-3’ and olMR7339, 5’-GTAGGTGGAAATTCTAGCATCATCC-3’ were used as internal PCR controls.

### Neuron cell culture

For obtaining embryonic hippocampal tissue, timed pregnant mice were sacrificed by cervical dislocation. Cell cultures were prepared from e17.5-18.5 mouse embryos as described previously ^29^. Briefly, tissue was dissected in Hank’s solution (Merck), incubated in 0.25% trypsin (Life Technologies) and 1 mg/ml DNAse (Roche) for 15 min at 37°C and dissociated into single cells by gentle pipetting. Cells were seeded on poly-D-lysine coated glass cover slips in DMEM (Invitrogen) with 10% FBS and PenStrep (both Gibco). 2 h after plating, the medium was replaced with Neurobasal medium with 0.6% Glucose, 2% B27, Glutamax (all Life Technologies) and PenStrep (Gibco) and cells were kept in a 37°C incubator with 5% CO_2_ and a humidified atmosphere. At 3 DIV, 1 μM cytosine arabinoside (Sigma) was added to the medium.

### Immunofluorescence microscopy and expansion microscopy

Cells were grown on poly-L-lysine- or poly-D-lysine-(neurons) coated coverslips and fixed with methanol at −20°C for a minimum of 15 min or with 3.7% paraformaldehyde at 37°C, followed by methanol at −20°C as described for the microtubule regrowth assays. To visualize centrioles with α-tubulin or acetylated α-tubulin, cells were incubated on ice for 30 min to depolymerize cytoplasmic microtubules before fixation. To remove cytoplasmic background (stainings of centrioles for α−tubulin or EGFP-HAUS6) cells were pre-extracted in ice-cold PHEM (60 mM PIPES, 25 mM HEPES, 10 mM EGTA, 2 mM MgCl_2_) pH 6.9 with 0.1% Triton X-100 for 1-2 min before fixation. Fixed cells were washed with PBS and blocked in PBS-BT (PBS, 3% BSA, 0.1% Triton X-100) for 1 h at RT, followed by the incubation with primary antibodies in PBS-BT either for 1 h at RT or overnight at 4°C. After washes in PBS-T (PBS, 0.1% Triton X-100) cells were incubated with secondary antibodies and 0.5 μg/ml DAPI (where appropriate) in PBS-BT for 1 h at RT. Cells were washed in PBS-T and either mounted in ProLong Gold Antifade (Thermofisher) on glass slides or further processed for ExM, as described previously ^51^: cells were washed with PBS and subsequently incubated in 0.1 mg/ml Acryloyl X (Life Technologies) in PBS at RT overnight. Cells were washed in monomer solution (1 x PBS, 2 M NaCl, 2.5% acrylamide (Bio-Rad), 0.15% Methylenbisacrylamide (Santa Cruz), 8.625% sodium acrylate (Sigma) and embedded in monomer solution containing 0.2% APS and 0.2% TEMED. Gels were polymerized for 2 h at 37°C and then digested with 8 U/ml Proteinase K (Invitrogen) in 50 mM Tris-HCl pH 8, 1 mM EDTA, 1 M NaCl, 0.5% Triton X-100 for 4 h at 37°C. Gels were expanded in MilliQ (expansion factor ~4) and mounted on poly-L-lysine-coated coverslips for imaging.

Images were acquired with an Orca AG camera (Hamamatsu) on a Leica DMI6000B microscope equipped with a 1.4 NA 100× oil immersion objective. AF6000 software (Leica) was used for image acquisition and blind deconvolution. Alternatively, images were acquired with an MRm camera on an Axiovert 200M (Carl Zeiss) using a 1.4 NA 63× Plan Apo objective and Axiovision software (Fig. 5c). Images were processed in ImageJ and Photoshop (Adobe) and represent maximum projections of a deconvolved stack or a single section. Image J was used for the quantification of fluorescence intensities. Images were acquired with constant exposure settings and pixel grey levels of the focused z plane were measured within a region of interest (ROI) encompassing a single centriole. Background fluorescence was measured adjacent to the ROI and subtracted.

### Western blotting (WB)

Cells were washed in PBS and lysed in 50 mM HEPES, pH 7.5, 150 mM NaCl, 1 mM MgCl_2_, 1 mM EGTA, 0.5% NP-40 and protease inhibitors (Roche) for at least 10 minutes on ice. Extract was cleared by centrifugation and subjected to SDS PAGE, followed by the transfer of proteins to PVDF membranes by tank blotting. Subsequently, membranes were blocked and probed with antibodies.

### Antibodies

Generation of rabbit polyclonal antibodies against HAUS6 (WB, 1:2000, IF, 1:1000 or 1:500 (ExM)) and GCP4 (ExM, 1:100) has been described previously ^7,52^. Other antibodies used in this study were: mouse anti-γ-tubulin (TU-30, Exbio; IF, 1:500 or 1:250 (ExM)), rabbit anti-γ-tubulin R75 ^53^ (1:1000), rabbit anti-α-tubulin (ab18251, Abcam; ExM, 1:300), mouse anti-acetylated α-tubulin (clone 6-11B-1, Merck; ExM, 1:250), mouse anti-acetylated α-tubulin (T7451, Sigma; IF, 1:1000), mouse anti-polyglutamylated tubulin (GT335, AdipoGen; ExM, 1:250), rabbit anti-NEDD1 ^4^ (ExM, 1:250), rabbit anti-HAUS5 ^19^ (ExM, 1:100), rabbit anti-pericentrin ^4^ (ExM, 1:250), rabbit anti-GFP (A6455, Invitrogen; ExM, 1:250), chicken anti-GFP (GFP-1020, Aves Labs; ExM, 1: 1:250), mouse anti-centrin 1 (clone 20H5, Millipore; IF, 1:500 or 1:250 (ExM)), rabbit anti-POC5 (A303-341A, Bethyl Laboratories; WB, 1: 2500, IF: 1:500 or 1:250 (ExM)), mouse anti-SAS-6 (sc-81431, Santa Cruz; IF, 1:100 or 1:50 (ExM)), mouse anti-centrobin ^54^ (IF, 1:500), rabbit anti-ninein ^55^ (IF, 1:100), rabbit anti-ODF2 (43840, Abcam; IF, 1:500), mouse anti-ARL13B (sc-515784, Santa Cruz; IF, 1:100), rabbit anti-CEP192 ^24^ (ExM, 1:500), mouse anti-GAPDH (sc-47724, Santa Cruz Biotechnology; WB, 1:10000). Alexa-Fluor-488-, Alexa-Fluor-568- and Alexa-Fluor-647-conjugated, cross-adsorbed secondary antibodies were obtained from Thermo Fisher (1:500 or 1:100 (ExM)). Streptavidin Alexa Fluor 594 was obtained from Invitrogen (1:5000). Horseradish-peroxidase-coupled secondary antibodies for WB were obtained from Jackson ImmunoResearch Laboratories (1:5000).

### Plasmids

The EGFP-HAUS8 expression plasmid was provided by Laurence Pelletier ^19^. The POC5-GFP expression plasmid was obtained from Ciaran Morrison ^30^. The plasmid expressing EGFP-HAUS6 was generated by cloning HAUS6 cDNA into pCS2-EGFP using AscI and Fse1 restriction sites. Site-directed mutagenesis was used to render EGFP-HAUS6 RNAi-insensitive (HAUS6 591A>G; 594T>C; 597G>A; 600G>C). A plasmid for expression of BirA-HAUS6 was generated by subcloning RNAi-insensitive HAUS6 into pCDNA5 FLAG-BirA^R118G^ (provided by Brian Raught) ^56^. Subsequently, FLAG-BirA^R118G^– HAUS6 was amplified by PCR and inserted into pEGFP-N1, replacing EGFP.

### Microtubule regrowth assay

U2OS cells were grown on poly-L-lysine-coated coverslips and incubated on ice for 40 min to depolymerize cytoplasmic microtubules. For microtubule regrowth, coverslips were transferred into 3.7% paraformaldehyde at 37°C for 1 min, followed by the incubation in methanol at −20°C for a minimum of 15 min. For the negative control (no microtubule regrowth), cells were incubated in 3.7% paraformaldehyde on ice for 10 min, followed by the incubation in methanol at −20°C for a minimum of 15 min.

### Transfection of plasmid and siRNA

Lipofectamine 2000 (Invitrogen) was used for the transfection of expression plasmids. Depletion of CEP192, HAUS6, NEDD1 and POC5 was performed by transfecting cells with the following siRNA oligonucleotides (Sigma) CEP192, 5’-AAGGAAGACAUUUUCAUCUCU-3’; HAUS6, 5’-CAGUUAAGCAGGUACGAA-3’; NEDD1, 5’-GCAGACAUGUGUCAAUUUGTT-3’; POC5, 5’-CAACAAAUUCUAGUCAUA-3’; using Lipofectamine RNAiMAX (Invitrogen). siRNA oligos against luciferase (5’-UCGAAGUAUUCCGCGUACG-3’) were used as control.

### Centrosome isolation and BioID

BioID from centrosomes was performed as described previously ^27^. Briefly, ten 15 cm dishes of U2OS cells stably expressing BirA-HAUS6 or parental U2OS cells (negative control) were incubated in culture medium containing 50 μM Biotin (Bio Basic) overnight. Subsequently, the medium was replaced and cells were incubated in culture medium containing 5 μg/ml nocodazole (Sigma) for 1 h. Cells were then washed with ice-cold HB buffer (20 mM HEPES pH 7.8, 5 mM K-acetate, 0.5 mM MgCl_2_, 0.5 mM DTT) and protease inhibitors (Roche) and incubated in 3 ml HB buffer for 10 min at 4°C. Cells were scraped from the plate and transferred into a 15 ml dounce homogenizer. After homogenization the lysate was centrifuged at 1500xg at 4°C for 5 min to pellet nuclei. Supernatant was collected and the pellet was washed again with HB buffer and centrifuged. Both supernatants were combined and 0.1% Triton X-100 was added, before the lysate was centrifuged at 1500xg at 4°C for 5 min. Supernatant was collected and loaded onto a sucrose gradient (discontinuous sucrose gradient prepared with 5 ml of 70%, 3 ml of 50% and 3 ml of 40% sucrose) in Ultra-Clear Beckman tubes and centrifuged at 26000 rpm (SW32Ti rotor, Beckman Coulter) for 1 h at 4°C. 500 μl fractions were collected and centrosome enrichment was determined by analyzing 10 μl of each fraction by WB using anti-γ-tubulin and anti-centrin antibodies. Centrosome-containing fractions were pooled and resuspended in lysis buffer (50 mM Tris pH 7.4, 500 mM NaCl, 0.4% SDS, 5 mM EDTA, 1 mM DTT, 2% Triton X-100, protease inhibitors). Samples were sonicated and equal volumes of ice-cold 50 mM Tris pH 7.4 was added. Lysates were centrifuged at 15000 rpm (JA 25.50 rotor, Beckman Coulter) for 10 min at 4°C. Supernatant was added to streptavidin agarose resin and incubated for 3 h at 4°C. Beads were washed several times with different wash buffers (buffer 1: 2% SDS in H_2_O; buffer 2: 0.2% deoxycholate, 1% Triton X-100, 500 mM NaCl, 1 mM EDTA, 50 mM HEPES pH 7.5; buffer 3: 10 mM Tris pH 8.1, 250 mM LiCl, 0.5% NP-40, 0.5% deoxycholate, 1% Triton X-100, 500 mM NaCl, 1 mM EDTA; buffer 4: 50 mM Tris pH 7.4, 50 mM NaCl) and finally resuspended in 100 μl of 50 mM NH_4_HCO_3_.

### Mass spectrometry analysis

Samples were digested with 1.08 μg (0.1 μg/μl) trypsin in 50 mM NH_4_HCO_3_ at 37°C overnight. Additional 1.08 μg trypsin was added and samples were incubated for 2 h at 37°C before formic acid was added (1% final concentration). Samples were cleaned through C18 tips (polyLC C18) and peptides were eluted with 80% acetonitrile/1% formic acid and diluted to 20% acetonitrile/0.25% formic acid before loading into strong cation exchange columns (polyLC SCX). Peptides were eluted in 5% NH_4_OH/30% methanol. Samples were evaporated to dryness and reconstituted in H_2_O with 3% acetonitrile/1% formic acid in a total volume of 50 μl. For mass spectrometry analysis, the reconstituted sample was further diluted 1:8 in H_2_O with 3% acetonitrile/1% formic acid. Samples were injected by triplicate (5μl per injection).

Sample was loaded at a flow rate of 15 μl/min on a 300 μm × 5 mm PepMap100, 5 μm, 100 A, C18 μ-precolumn using a Thermo Scientific Dionex Ultimate 3000 chromatographic system (Thermo Scientific). Peptide separation was done with a 90 min run on a C18 analytical column (Acclaim PepMapR RSLC 75 μm × 50 cm, nanoViper, C18, 2 μm, 100 A, Thermo Scientific), comprising three consecutive steps with linear gradients from 3 to 35% B in 60 min, from 35 to 50% B in 5 min, and from 50% to 85% B in 2 min. Isocratic elution was done at 85% B in 5 min and stabilization to initial conditions (A= 0.1% formic acid in H_2_O, B= 0.1% formic acid in CH_3_CN). The outlet of the column was directly connected to a TriVersa NanoMate (Advion) fitted on an Orbitrap Fusion Lumos™ Tribrid (Thermo Scientific). The mass spectrometer was operated in a data-dependent acquisition mode, survey MS scans were acquired with a resolution of 120,000 (defined at 200 m/z), and lock mass was defined at 445.12 m/z in each scan. In each scan the top speed (most intense) ions were fragmented by CID and detected in the linear ion trap. The ion count target values for survey and MS/MS scans were 400,000 and 10,000, respectively. Target ions already selected for MS/MS were dynamically excluded for 15 s. Spray voltage in the NanoMate source was set to 1.60 kV. RF Lens was tuned to 30%. Minimal signal required to trigger MS to MS/MS switch was set to 5,000. The spectrometer was working in positive polarity mode and singly charge state precursors were rejected for fragmentation. Database searching was done with Proteome Discoverer software v2.1.0.81 (Thermo) using Sequest HT search engine and SwissProt Human release 2018 01 and manually introduced contaminants database and user proteins. Searches against targeted and decoy database were used for determining the false discovery rate (FDR). Search parameters for trypsin enzyme specificity allowed for two missed cleavage sites, oxidation in M and acetylation in protein N-terminus. Peptide mass tolerance was 10 ppm and the MS/MS tolerance was 0.6 Da. Peptides with q-value lower than 0.1 and FDR < 1% were considered as positive with a high confidence level.

For the quantitative analysis contaminant identifications were removed and unique peptide spectrum matches of protein groups identified with Sequest HT were analyzed with SAINTexpress-spc v3.11 ^57^. High confidence interactors were defined as those with Bayesian false discovery rate *BFDR* ≤ 0.02.

### Statistical analysis and replication of experiments

Statistical analysis was performed using Prism 7 software. For quantifications of the accumulation of HAUS6/NEDD1 at distinct centriolar sites (Fig. 1f), centriole numbers (Fig. 4b,d,e,g, Supplementary Fig. 4e), the mean of independent experiments was first determined and statistics were performed on the entirety of the obtained means. Normality of data distribution within a data set was tested with a D'Agostino-Pearson normality test (Fig. 1g, Fig. 5e, Supplementary Fig. 2c) or a Shapiro-Wilk normality test (Fig. 1f, Fig. 4b,d,e,g, Fig. 5. b,d, Supplementary Fig. 4c), depending on sample size. Significances between two data sets were determined using an unpaired two-tailed Student’s t-test (Fig. 1f,g, Fig. 4b,d,e,g, Fig. 5b,d,e, Supplementary Fig. 4c,e) or a Mann-Whitney test (Fig. 4e (data set with BI2563), Supplementary Fig. 2c). The results, together with the number of independent experiments and sample sizes, are reported in the figures and figure legends. Protein localizations/dependencies and microtubule regrowth along the centriole wall has been confirmed in at least two independent experiments, except for CEP192 wall localization and its loss upon CEP192 RNAi (Fig. 2b). The displayed images are representative examples. The BioID (Fig. 3a, Supplementary Fig. 3b) was performed once.

## Acknowledgements

NS was supported by an EMBO long-term fellowship (ALTF 820-2015) and a Marie Skłodowska-Curie Action fellowship: This project has received funding from the European Union’s Horizon 2020 research and innovation programme under the Marie Sklodowska-Curie grant agreement No 703907. ID was funded by the European Union's Horizon 2020 research and innovation programme under the Marie Skłodowska-Curie grant agreement No. 754510. LH and AM were in part supported by grant 13-BSV8-0007-01 from “Agence Nationale de la Recherche” (France), and by grant SFI20121205511 from “Fondation ARC pour la recherche sur le cancer”. JL acknowledges support by grants BFU2015-69275-P (MINECO/FEDER), PGC2018-099562-B-I00 (MICINN), 2017 SGR 1089 (AGAUR) and by intramural funds of IRB Barcelona, recipient of a Severo Ochoa Centre of Excellence Award from the Spanish Ministry of Science and Innovation and supported by CERCA (Generalitat de Catalunya). We thank the IRB Mass Spectrometry & Proteomics Core Facility, a member of ProteoRed, PRB3-ISCIII, supported by grant PRB3 (IPT17/0019-ISCIIISGEFI/ERDF), and the Animal Facility of the Parc Scientific de Barcelona for excellent support. We thank Hélène Dollfus (University of Strasbourg, Strasbourg, France), Meng-Fu Bryan Tsou (Memorial Sloan Kettering Cancer Center, New York, USA), Laurence Pelletier (Lunenfeld-Tanenbaum Research Institute, Toronto, Canada), Ciaran Morrison (National University of Ireland, Galway, Ireland) and Brian Raught (Ontario Cancer Institute and Department of Medical Biophysics, University of Toronto, Toronto, Canada) for cells and reagents.

## Author contributions

NS designed experimental strategies, performed most of the experiments, prepared figures and contributed to manuscript writing. LH performed experiments with patient fibroblasts. RV generated conditional *Haus6* KO mice and prepared neuronal cultures. CL supervised animal experiments and performed the BioID experiment. ID performed some of the POC5 RNAi experiments in RPE1 *p53 -/-* cells. AM supervised experiments with patient fibroblasts. JL supervised the study, proposed experimental strategies, and contributed to manuscript writing.

---

**Supplementary Figure 1.**
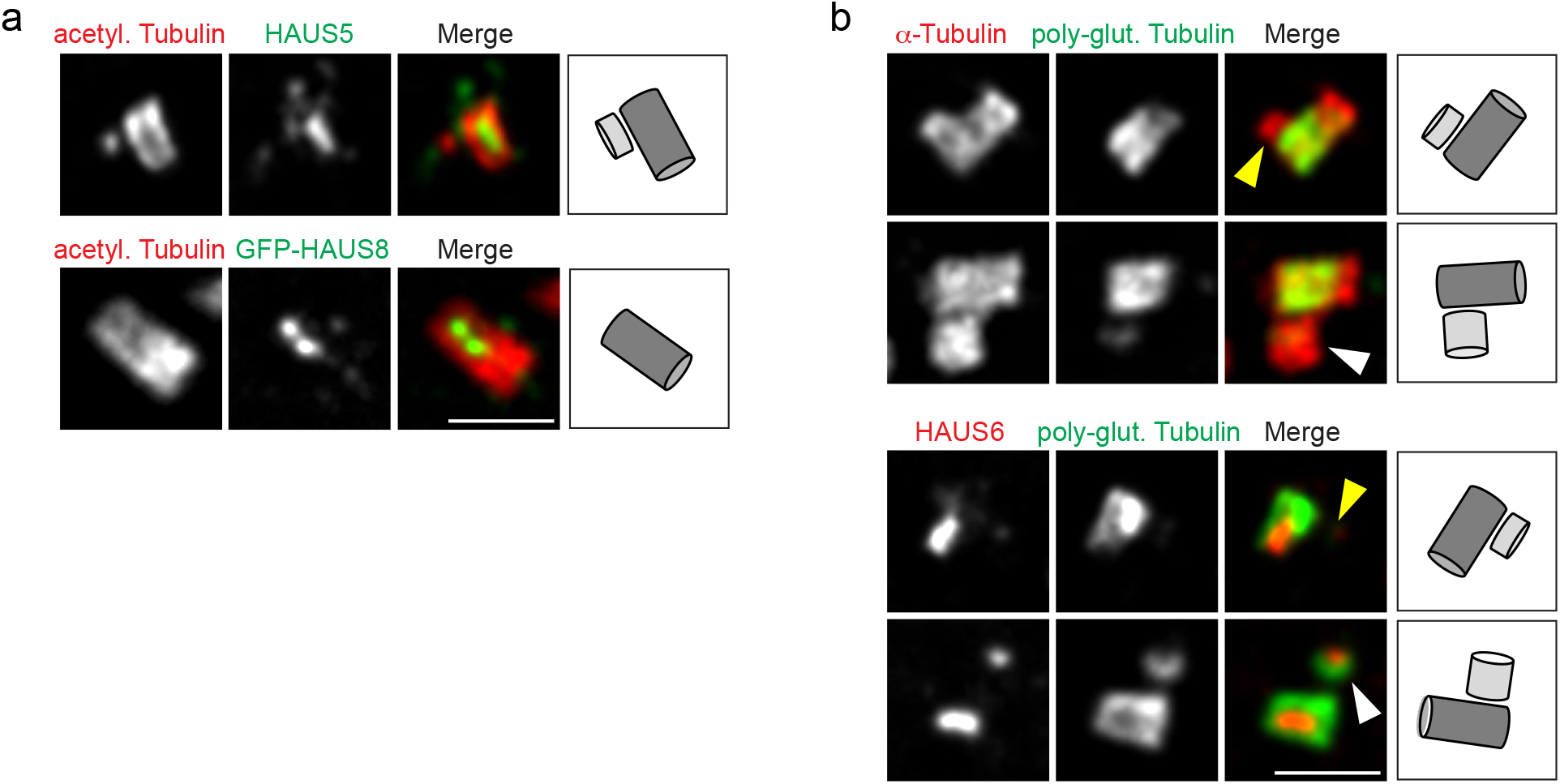
Augmin localizes to the centriole lumen late during the cell cycle. (**a**) Centrioles of parental U2OS cells or U2OS cells stably expressing EGFP-HAUS8 in ExM stained for acetylated α-tubulin (red) and HAUS5 (green) or GFP (EGFP-HAUS8, green). (**b**) Centrioles of U2OS cells in ExM stained for α-tubulin (red) and poly-glutamylated tubulin (green) or HAUS6 (red) and poly-glutamylated tubulin (green). Yellow arrowheads point to daughter centrioles that lack HAUS6/poly-glutamylation, white arrowheads point to poly-glutamylated daughter centrioles (that have HAUS6 in the lumen). Bar (all panels), 2 μm. Cartoons illustrate centriole configurations in the corresponding panels, dark grey = mother centriole, light grey = daughter centriole.

**Supplementary Figure 2.**
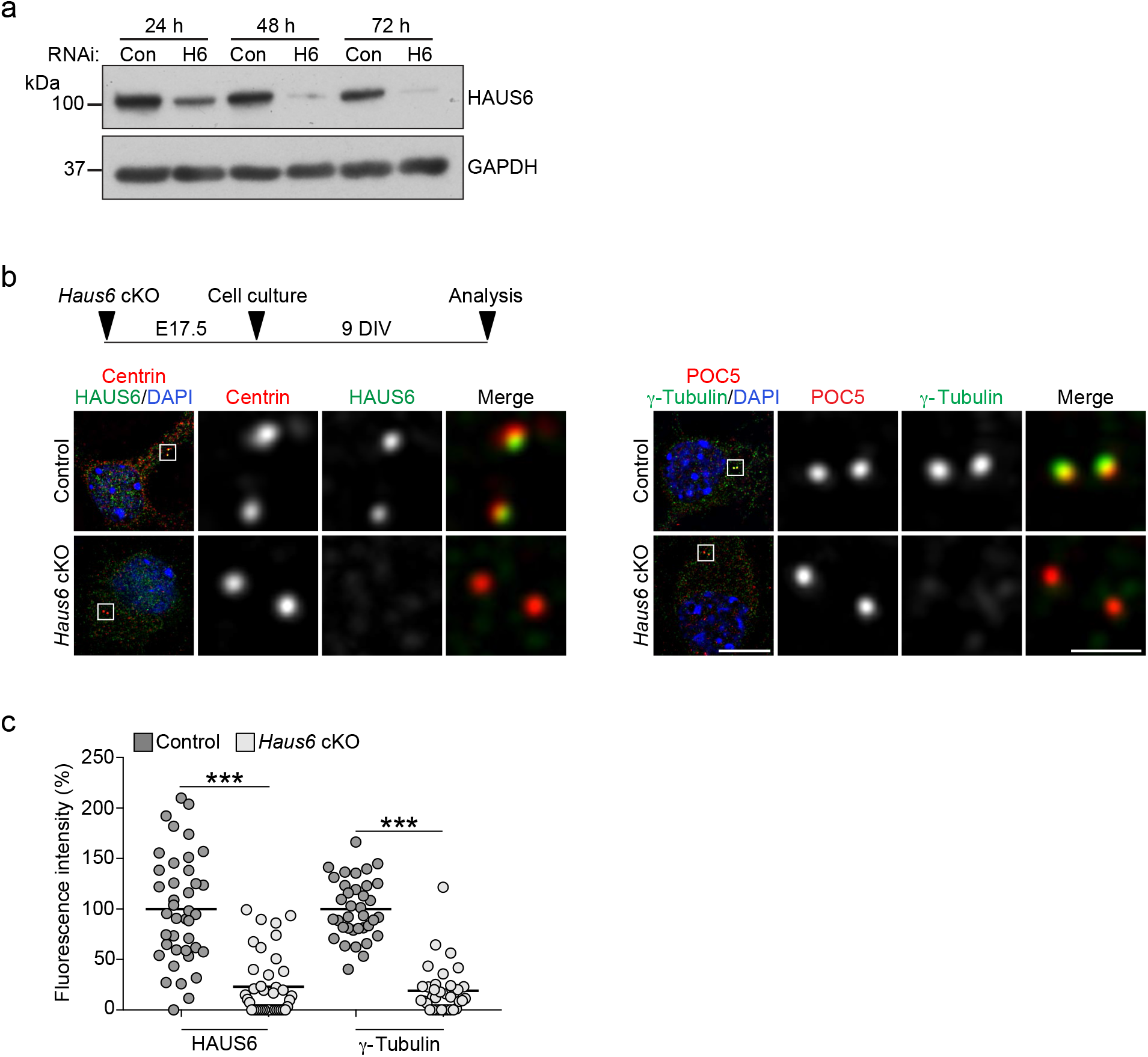
γTuRC centriole lumen localization depends on augmin. (**a**) Western Blot analysis of HAUS6 from control RNAi and HAUS6 RNAi U2OS cells at different time points after siRNA transfection. GAPDH was used as loading control. (**b**) Murine control and *Haus6* cKO neurons at 9 DIV stained for centrin (red), HAUS6 (green) and DNA (DAPI, blue) or POC5 (red), γ-tubulin (green) and DNA (DAPI, blue). Bar, 10 μm (merge with DAPI) or 1 μm (insets depicting centrioles). The experimental design is depicted schematically. (c) Fluorescence signals for HAUS6 and γ-tubulin at centrioles in control and *Haus6* cKO neurons at 9 DIV, plotted as percentages of the signal in control cells (average intensity in control cells was set to 100%). The mean is depicted as horizontal line (HAUS6 fluorescence intensity in control and *Haus6* cKO, 100% ± 53.4% and 22.9% ± 30.8%, mean ± SD, n = 40 centrioles from 20 cells per condition, combined from two independent experiments, p < 0.001; γ-tubulin fluorescence intensity in control and *Haus6* cKO, 100% ± 29.2% and 19.0% ± 23.8%, mean ± SD, n = 36 centrioles from 18 cells per condition, combined from two independent experiments, p < 0.001). *** p < 0.001.

**Supplementary Figure 3.**
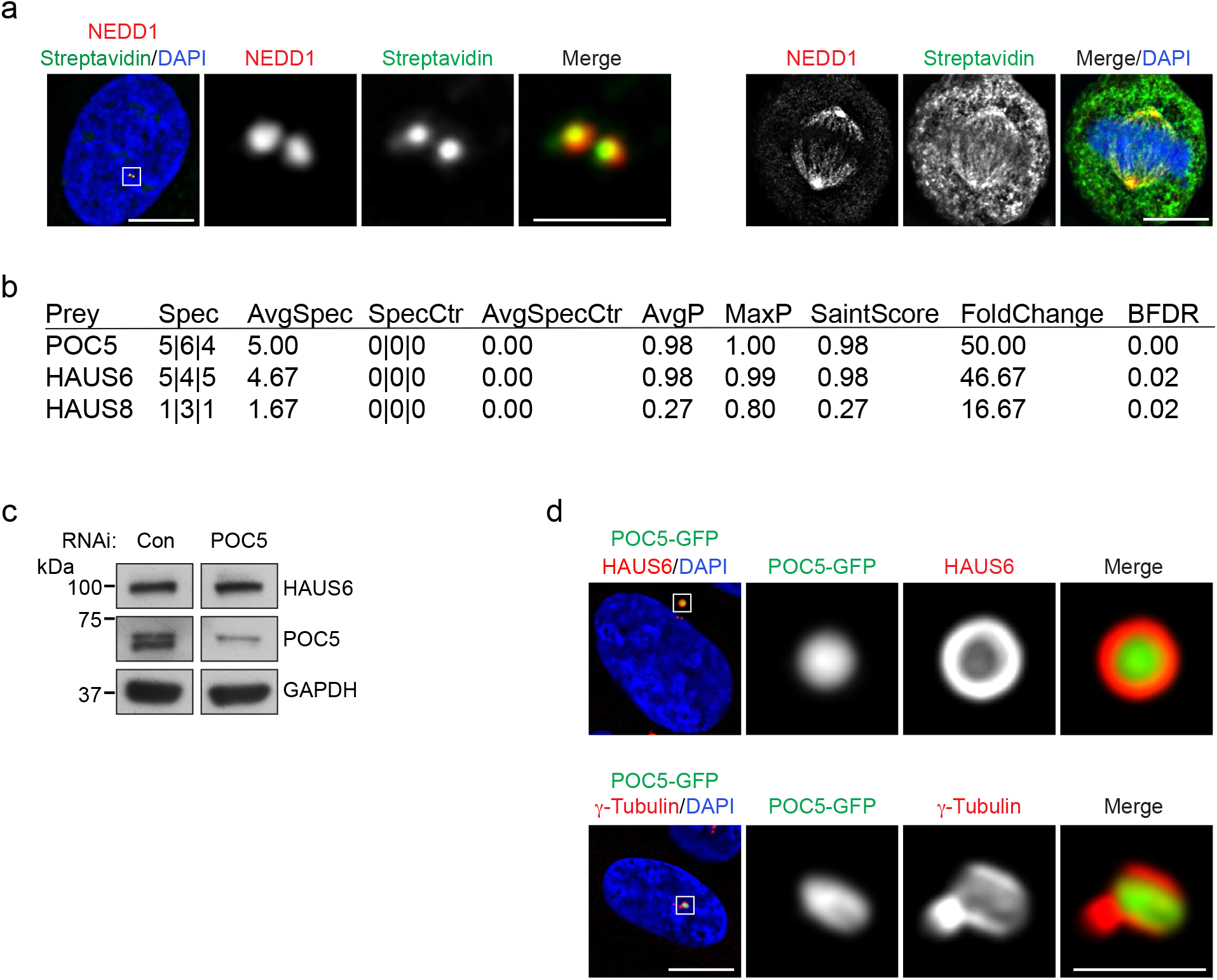
Identification of POC5 as a proximity interactor of HAUS6. (**a**) Interphase or mitotic U2OS cells stably expressing BirA-HAUS6 in the presence of biotin stained for NEDD1 (green), biotin (568-conjugated streptavidin, red) and DNA (DAPI, blue). (**b**) Mass spectrometry analysis of proximity interactors of BirA-HAUS6 from isolated centrosomes of U2OS cells. (**c**) Western Blot analysis of POC5 from control RNAi and POC5 RNAi U2OS cells. GAPDH was used as loading control. (**d**) U2OS cells stably expressing POC5-GFP stained for GFP (POC5-GFP, green), HAUS6 (red) or γ-tubulin (red) and DNA (DAPI, blue). Bar (all panels), 10 μm or 2 μm (insets).

**Supplementary Figure 4.**
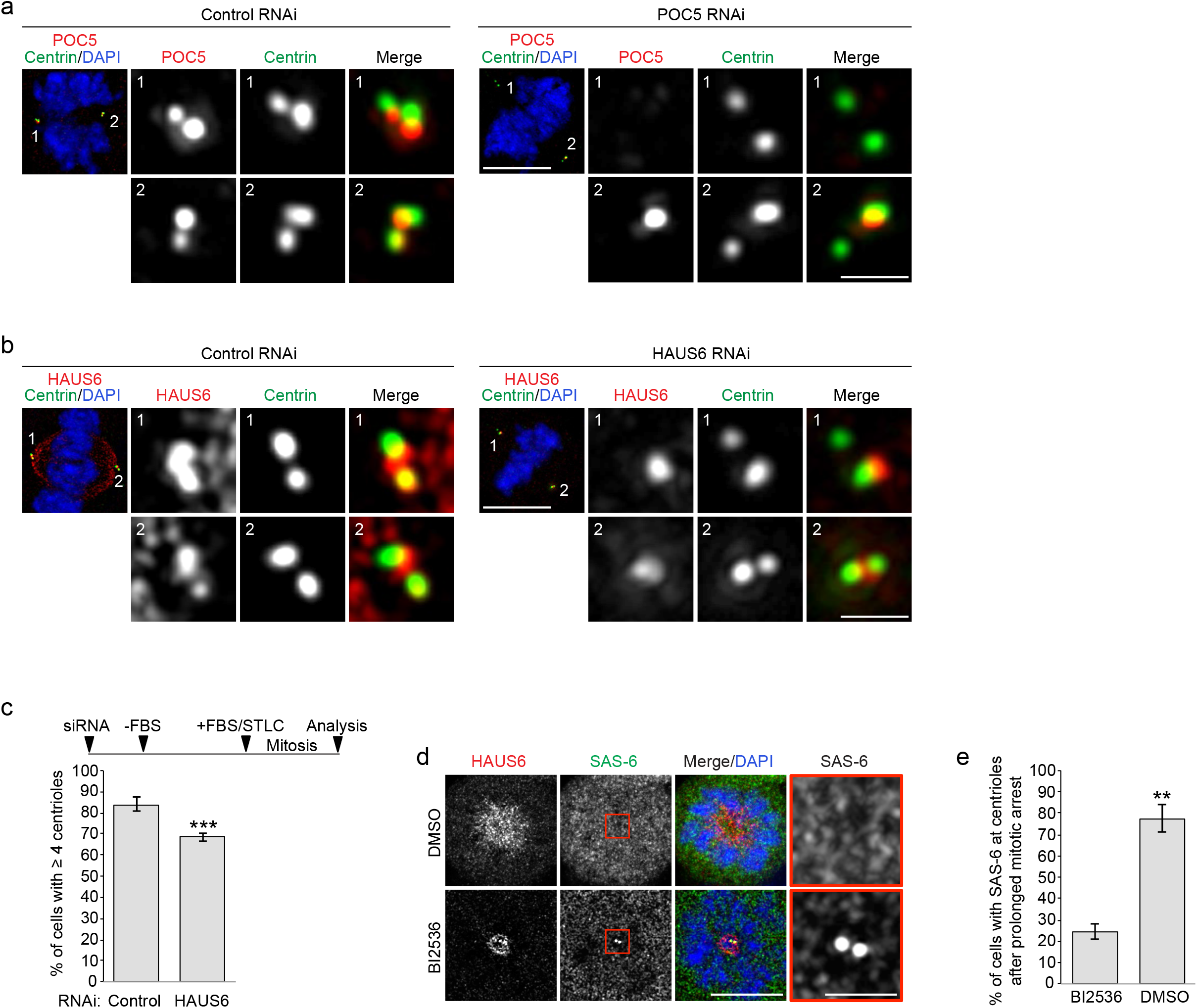
Centriole numbers are not reduced in cycling POC5 RNAi or HAUS6 RNAi cells. (**a**) Mitotic control RNAi or POC5 RNAi U2OS cells stained for POC5 (red), centrin (green) and DNA (DAPI, blue). (**b**) Mitotic control RNAi or HAUS6 RNAi U2OS cells stained for HAUS6 (red), centrin (green) and DNA (DAPI, blue). (**c**) Quantifications of the percentage of control RNAi or HAUS6 RNAi hTERT RPE1 cells with ≥ 4 centrioles (centrin foci) after prolonged mitotic arrest. Error bars represent standard deviations from the mean obtained from four independent experiments (control RNAi: 84.0 ± 3.2%, HAUS6 RNAi: 68.5 ± 1.9%; mean ± SD, 100-500 cells per condition and experiment, p < 0.001). The experimental design is depicted schematically. Cells were transfected with siRNA oligos and 5 h later serum-starved for ~73 h. Subsequently, cells were released into medium containing FBS and STLC for ~25 h before cells were fixed. Control RNAi U2OS cells which spent 24 h in mitosis, with (BI2536) or without (DMSO) PLK1 inhibition, stained for HAUS6 (red), SAS-6 (green) and DNA (DAPI, blue). (**e**) Quantifications of the percentage of cells with SAS-6 at centrioles, with (BI2536) or without (DMSO) PLK1 inhibition after spending 24 h in mitosis. Error bars represent standard deviations from the mean obtained from two independent experiments (without PLK1 inhibition: 24.5 ± 3.5%, with PLK1 inhibition: 77.5 ± 6.4%, mean ± SD, 100-200 cells per condition and experiment, p < 0.01). ** p < 0.01. Bar (all panels), 10 μm or 1 μm (insets).

